# A new causality measure

**DOI:** 10.1101/446567

**Authors:** Kaushik Majumdar, Puneet Dheer

## Abstract

Let there are *n* i.i.d observations, each producing a two tuple of real values (*x_i_*, *y_i_*), *i* ∊ {1,……, *n*} giving rise to random variables *X =* (*x*_1_,…, *x_n_*) and *Y =* (*y*_1_,…, *y_n_*). What is the statistical significance of the hypothesis that an increase in the independent variable *X* is causing an increase (or decrease) to the dependent variable *Y* ?

## 1. Introduction

Complex partial seizures impair consciousness of the patient. Impairment of consciousness can be scored (Arthuis et al, 2009). The more is the score the higher is the loss of consciousness. Seizure is caused by abnormally high synchronization in the brain. Synchronization among brain electrical signals, collected through multiple electrodes, can be measured in many different ways. Almost all of them become high during the seizure compared to before or after it. However, there is no linear or straightforward relationship between the value of synchronization and consciousness score. There may be a small number of seizures recorded from a few patients with sufficient duration to study the effects on consciousness. How to establish a causal relationship between the synchronization values and consciousness scores of the seizures? One question of clinical importance is “Does enhanced synchronization lead to enhanced consciousness score leading to loss of consciousness?” If *X* gives synchronization values and *Y* consciousness scores, the values in *X* and *Y* can appear in any order and therefore the conventional causality measures like Granger causality will not be applicable. Cross correlation or chi-square independence will also not be applicable and will not be able to tell if enhanced synchronization is causing loss of consciousness or enhanced loss of consciousness is causing enhanced synchronization. Even for high cross correlation under certain ordering the causality can be statistically insignificant.

## 2. Proposed Method

### Case I: All the values of *X* are different

We are interested in the hypothesis that one particular trend in *X* (independent variable) is causing one particular trend in *Y* (dependent variable). Since values of observations are not in any particular order, let us increasingly order the values of *X*. Let *X^o^* = (*x*_*i*_1__, *x*_*i*_2__, *x*_*i*_3__,…, *x*_*i*_*n*__) is the new (increasing) order of *X*, whereas *X^o^* indicates the ordered *X*. So, *Y^o^* = (*y*_*i*_1__,*y*_*i*_2__,*y*_*i*_3__,…*y*_*i*_*n*__) indicates ordered *Y* but not in increasing or decreasing sense. If increasing *X* causes increasing (decreasing) *Y*, then *Y^o^* should be in increasing (decreasing) order. Particularly, decreasing (increasing) *Y^o^* will support the null hypothesis.?

Let us consider the difference operation 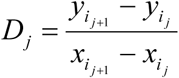, whose denominator is always positive and therefore sign of *D_j_* depends on the sign of the numerator. If the alternative hypothesis *H*_1_ states that, increase in *X* will cause increase in *Y*, then *D_j_* < 0, *j* ε {1,…,n −1} will support the null hypothesis *H*_0_. Let *D_j_* < 0 for *k* different values of *j* out of a total of *n* −1 values. What is the cumulative probability that *k* number of *D_j_* s will be negative?

Let us allow the odd, i.e. *D_j_* <0, to take a ‘fair’ probability of 1/ 2, which implies the probability of *D_j_* ≥0 is also 1/2. The cumulative probability that *k* number of *D_j_* s will be negative is therefore going to be a binomial distribution measure 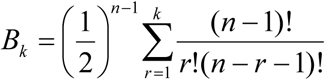. If *B_k_* < *α*, where *α* is the acceptable level of significance, we can say that *H_1_* is true with statistical significance *p* < *α*.

### Case II: Not all values of *X* are different

Let 1 ≤ *i* < *j* ≤ *n* and *x*_*i*_ = *x*_*j*_·. If *y*_*i*_ = *y*_*j*_ we can simply remove the *i* th or the *j* th observation from the analysis and go ahead with the remaining *n* −1 observations.

If *y_i_* ≠ *y*_*j*_, form two sets, each containing *n* −1 observations. One of them will contain the *i* th, but not the *j* th observation and the other contains the *j* th, but not the *i* th observation. Continue to divide the set of *n* observations unless each subset of observations satisfies the condition of Case I. Let there are *d* number of reduced sets. If we are looking for a p-value 0 < *α <* 1, then each of the *d* sets will have to have p-value for accepting *H*_1_ as *α/d* by the unweighted Bonferroni criterion (Shaffer 1995).

## 3. Results

We have consciousness score of the 11 patients during 22 complex partial seizures recorded in them. The intensity of seizure has been measured by the amount of synchronization it induced across all the seizure onset zone channels. We are interested in the hypothesis – *increment in the synchronization index increases the consciousness score*. Synchronization arranged in increasing order versus consciousness score has been plotted in Fig. 1. Statistical significance of the hypothesis – increased synchronization is increasing the consciousness score, is 0.0946 < 0.1. Table 1 shows the cross correlation between abscissa and ordinate of Fig. 1 is 0.44 with p = 0.041.

**Fig. 1.**
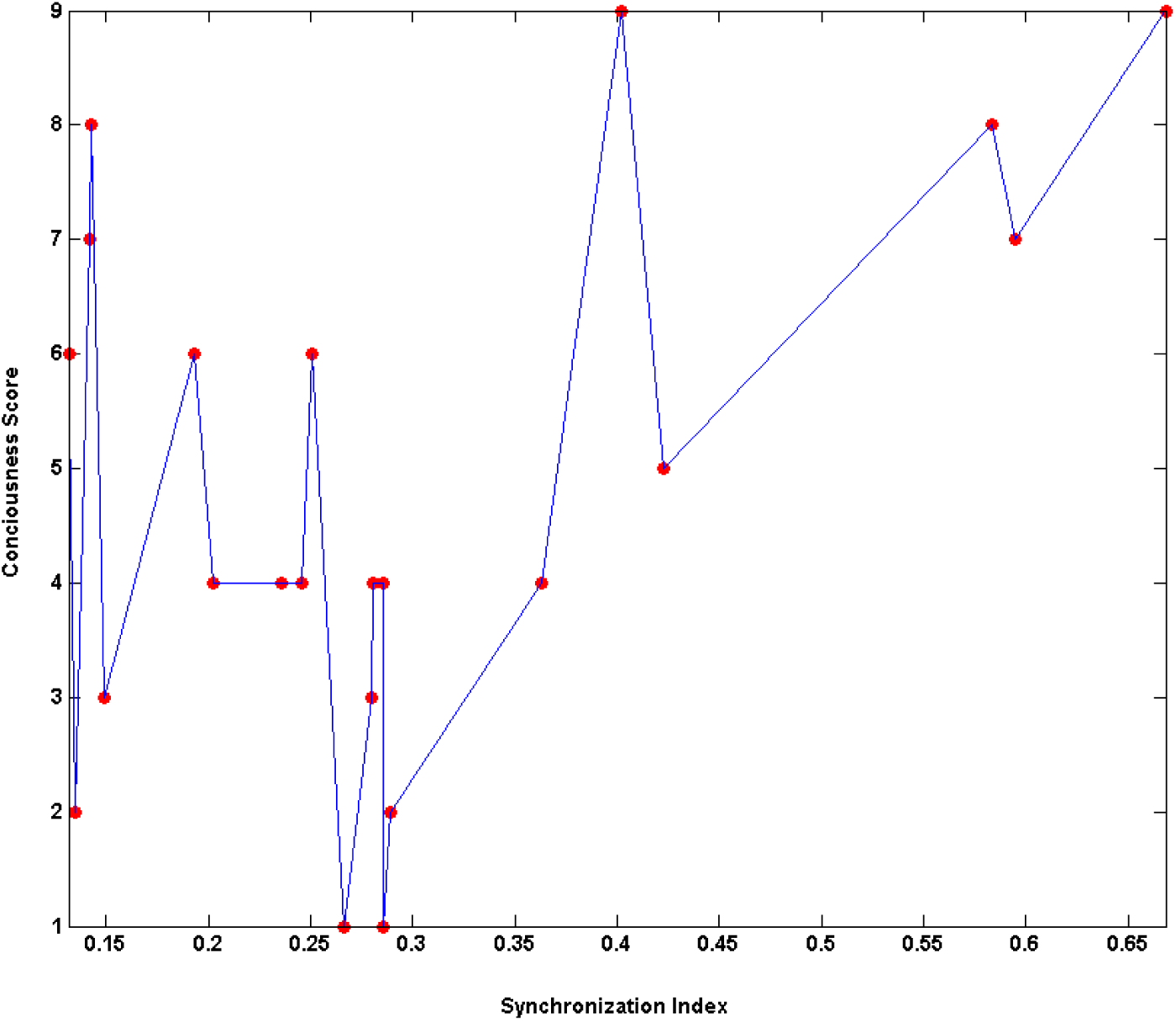
Synchronization index (mutual information across all the seizure onset zone channels in 0 to 1 scale) arranged in increasing order versus consciousness score (in 1 to 9 scale) plot of 22 complex partial seizures recorded in eleven patients.

**Table 1:** Correlations

|  |  | SI | CS |
| --- | --- | --- | --- |
| SI | Pearson Correlation | 1 | 0.440* |
|  | Sig. (2-tailed) |  | 0.041 |
|  | N | 22 | 22 |
| CS | Pearson Correlation | 0.440* | 1 |
|  | Sig. (2-tailed) | 0.041 |  |
|  | N | 22 | 22 |

\*Correlation is significant at the 0.05 level (2-tailed). Sig. = significance.

Now we will concentrate on the other side of the story – what is the statistical significance of the hypothesis that the increased consciousness score is driving the synchronization index up? Fig. 2 shows the two dimensional plot of synchronization index (along ordinate) of the 22 complex partial seizures recorded in 11 patients versus the consciousness score sorted in increasing order (abscissa). There are multiple synchronization index values associated with most of the single consciousness score values. The function mapping the consciousness score values onto the synchronization index is not a single valued function. There are 1152 different ways single valued functions can be constructed mapping the set of consciousness scores onto the set of synchronization indices. Fig. 3 shows the plot of one such function. In Fig. 3 the correlation between the nine consciousness scores and nine synchronization indices is 0.885082 with *p* = 0.001511, which is the lowest among all the 1152 cases (Fig. 4). Since in Fig. 3 the correlation between the abscissa and ordinate is the most significant and also quite high (0.885082), we have chosen this particular case to examine how statistically significantly the increment in the consciousness score is causing the increment in the corresponding synchronization indices. In Fig. 3 there are three negative *D_j_* s and five positive out of a total eight. The probability of *H*_0_ being true is the cumulative probability of up to three negative *D_j_* s occur in a total eight, which is 0.3633. This clearly shows even for high correlation value causality can be insignificant. Even if all eight are positive the p-value will be 0.0390625. Our predetermined acceptable p-value was 0.1. After the unweighted Bonferroni correction (Shaffer, 1995) the acceptable p-value for the hypothesis – increment in consciousness scores is causing increment in the corresponding synchronization indices, becomes 0.1/1152 = 0.00008681. It is clear that none of the 1152 cases will be able to attain this p-value to accept the hypothesis – increment in consciousness scores is causing increment in the corresponding synchronization indices.

**Fig. 2.**
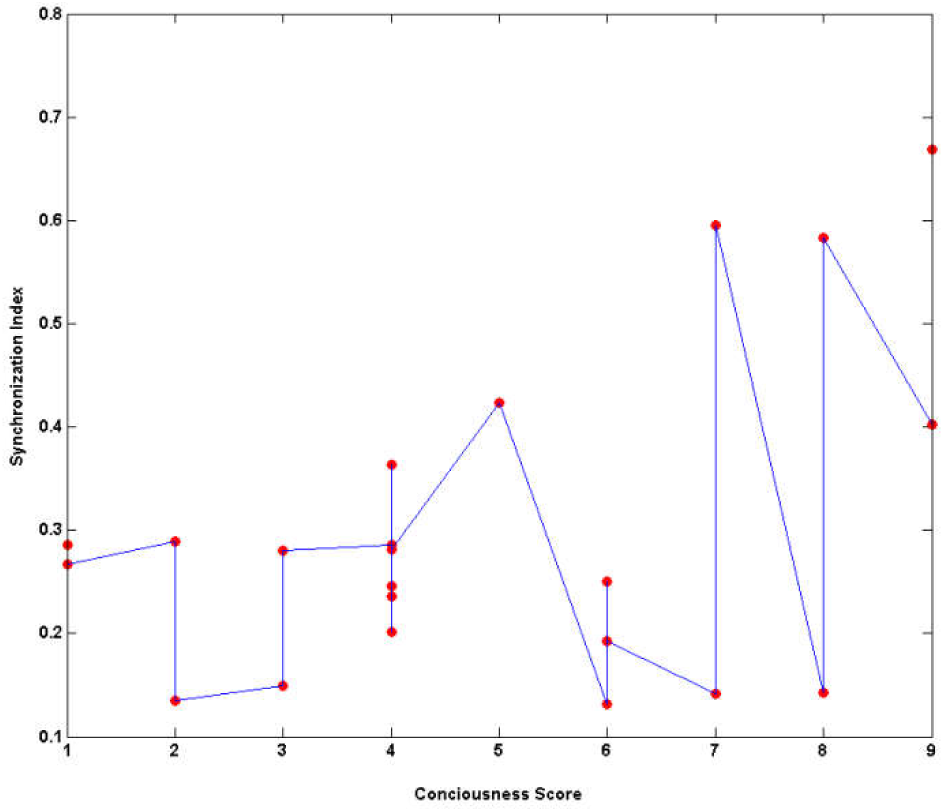
Consciousness score in increasing order makes the abscissa, whereas synchronization index makes the ordinate for the 22 complex partial seizure in Fig. 1.

**Fig. 3.**
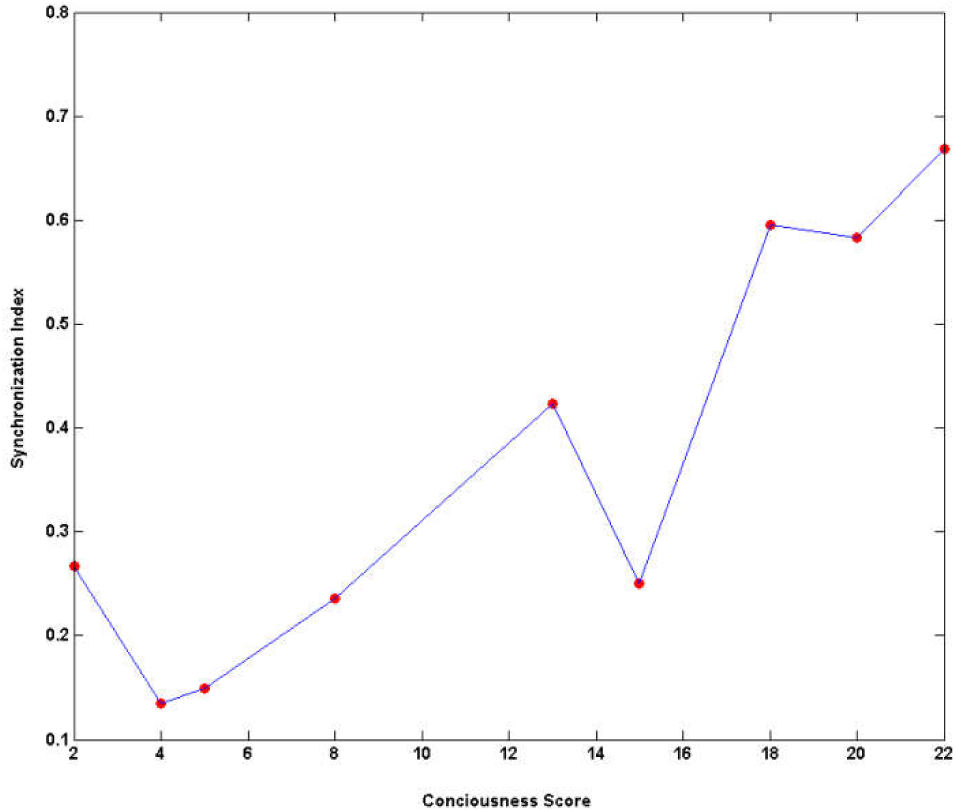
Sorted consciousness score in increasing order (abscissa) versus synchronization index (ordinate) plot with three negative *D.* values out of eight.

**Fig. 4.**
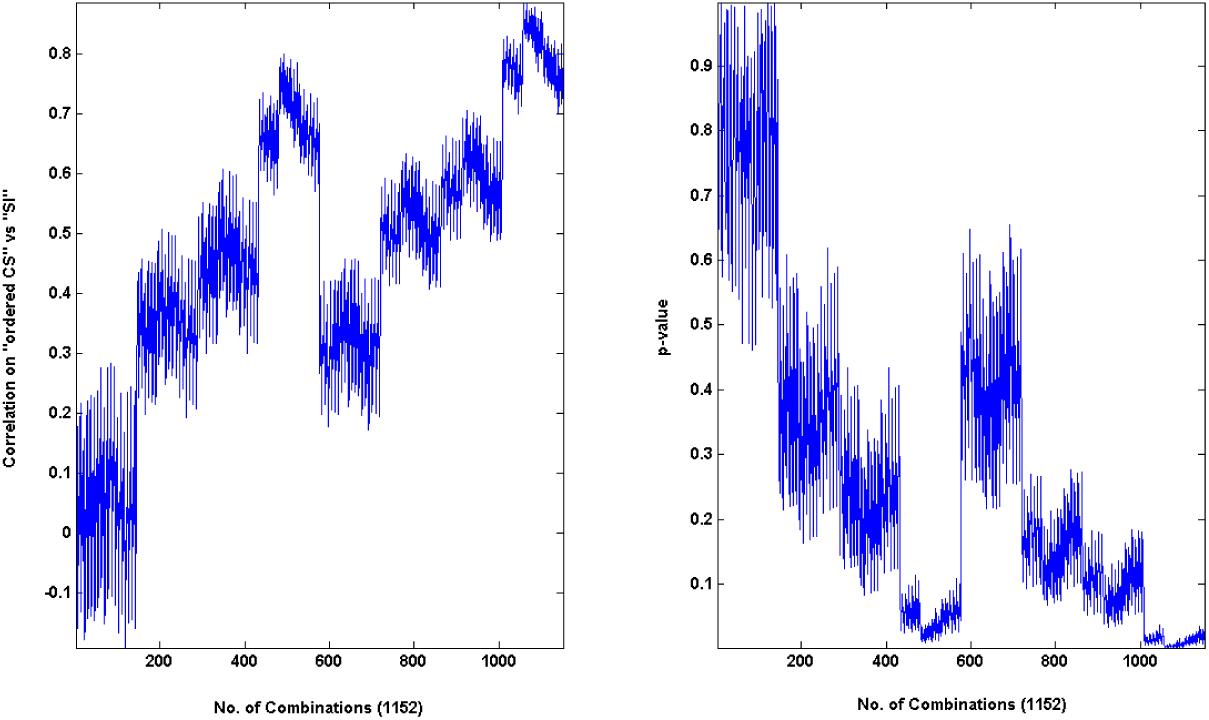
(Left) Plot of cross correlation values between the nine sorted consciousness scores in increasing order and the corresponding nine synchronization indices for all the 1152 cases. (Right) The corresponding p-values of all the 1152 cross correlation values in the left plot.

So, we conclude – increase in synchronization is causing increase in consciousness score, but not the other way round. In case increase in the independent variable causes decrease in the dependent variable, we have to take positive *D_j_* s as the *H*_0_ supporting events.

## Conclusion

When the independent variable can take one particular value multiple times, the Bonferroni correction may be too stringent a condition for the simultaneous multiple hypothesis testing. Under certain conditions relatively less stringent p-value may be chosen. For example, let 2*k* + *s* < *n* be true, where k is the number of signs of *D_j_* s favoring *H*_0_ in both the original set and the reduced set, *n* − *s* is the cardinality of the reduced set and *n* is the number of observations. It can be shown that the probability of *H*_0_ be true in the original set is bounded above by the probability of happening the same in the reduced set. So, the highest probability of *H*_0_ be true in all the reduced sets is an upper bound for the highest probability of *H*_0_ be true in the original set of observations. But this approach may not be very convenient from application point of view for the following reasons: (1) Number of reduced sets may be quite high (for example, the case we considered here it is 1152 for only 22 observations). (2) The number of signs of *D_j_* s favoring *H*_0_ may be unequal in the original and reduced sets of observations. When it is more in a reduced set then the probability of *H*_0_ being true in the original set cannot be bounded above by the probability of *H*_0_ being true in the reduced set.

*D_j_* can be negative, positive or zero for any *j* ∊ {1,……, *n* − 1}. We have assigned probability ½ to the event in which *D_j_* is taking sign favorable to *H*_0_. A data dependent more realistic probability measure for the event, in which *D_j_* is taking sign favorable to *H*_0_, will of course make the hypothesis testing more realistic.

## Acknowledgement

The work was supported by an Indian Statistical Institute grant no. SSIU-18/Syn/KM-New. Consciousness score was assigned to the patients by Sandipan Pati (Epileptologist) of University of Alabama at Birmingham and the synchronization indices were measured on the data provided by him.

